# NRF2 pathway activation reverts high-glucose-induced transcriptional memory in endothelial cells

**DOI:** 10.1101/2023.09.11.557207

**Authors:** Martí Wilson-Verdugo, Brandon Bustos-García, Olga Adame-Guerrero, Jaqueline Hersch-González, Nallely Cano-Dominguez, Maribel Soto-Nava, Carlos A. Acosta, Teresa Tusie-Luna, Santiago Avila-Rios, Lilia G. Noriega, Victor J. Valdes

**Affiliations:** Departamento de Biología Celular y del Desarrollo, Instituto de Fisiología Celular, Universidad Nacional Autónoma de México (UNAM), Ciudad de México. 04510, México; Centre for Research in Infectious Diseases of the National Institute of Respiratory Diseases (CIENI/INER), Mexico City 14080, Mexico; Hospital Diomed. Av Observatorio 354. Miguel Hidalgo. Ciudad de México, 11810. Mexico; Unidad de Biología Molecular y Medicina Genómica Instituto de Investigaciones Biomédicas UNAM / Instituto Nacional de Ciencias Médicas y Nutrición Salvador Zubiran, Ciudad de México, Mexico; Departamento de Fisiología de la Nutrición, Instituto Nacional de Ciencias Médicas y Nutrición Salvador Zubirán, Ciudad de México, Mexico

**Keywords:** Glucose, endothelial cells, metabolic memory, sulforaphane, NRF2, ATAC-seq, epigenetics, diabetes

## Abstract

Various diabetes complications, including nephropathy, retinopathy, and cardiovascular disease, arise from vascular dysfunction. In this context, it has been observed that past hyperglycaemic events can induce long-lasting transcriptional changes, a phenomenon termed “metabolic memory”. Yet, the underlying mechanisms driving these persistent effects are not fully characterized. In this study, we evaluated the genome-wide gene expression and chromatin accessibility alterations caused by transient high glucose exposure in human endothelial cells (ECs). We found that cells exposed to a transient high glucose episode had decreased glycolytic and oxygen consumption rates. Transcriptional profiling indicated that high glucose exposure induced substantial changes in the expression of genes belonging to pathways known to be impaired in diabetes, such as TGF-beta, TNF, FoxO, p53, and NRF2 pathways, many of which were retained after normalization of glucose concentrations. Furthermore, analysis of chromatin accessibility showed that transient hyperglycaemia can induce persistent modifications in the accessibility landscape, with the majority of differentially accessible regions located in non-promoter regions. Some of these regions were identified as putative enhancers with neighbouring genes persistently altered after transient high glucose exposure. Finally, we showed that activation of the NRF2 pathway through either NRF2 overexpression or supplementation with the plant-derived compound sulforaphane, was able to substantially revert the glucose-induced transcriptional memory in ECs. Our findings demonstrate that transient high glucose can induce persistent changes in both the transcriptomic and chromatin accessibility profiles of ECs, and that pharmacological NRF2 pathway activation is able to prevent and revert the high-glucose-induced transcriptional memory.

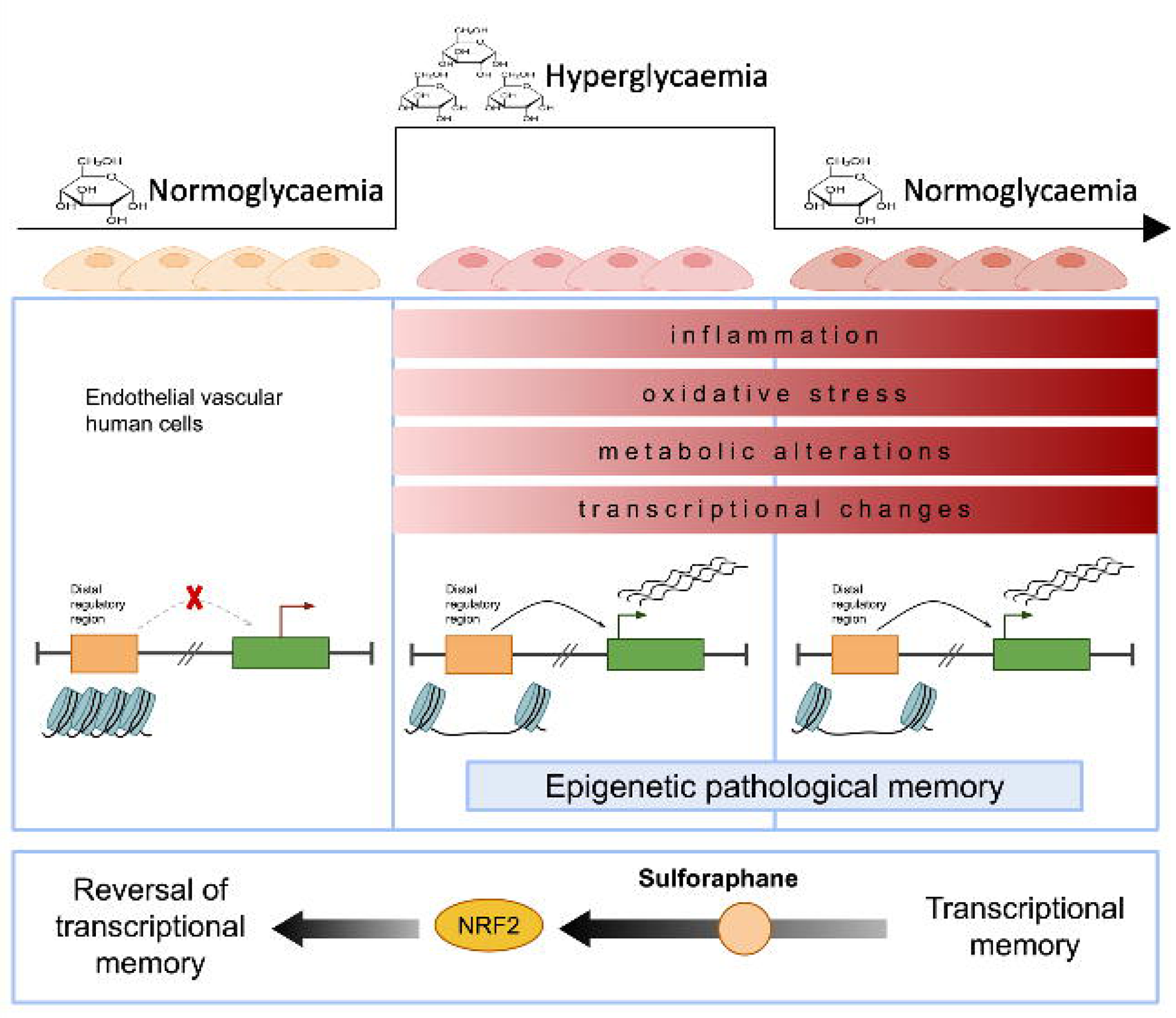

## 1. Introduction

Several diabetes-related pathologies, including nephropathy, retinopathy, and cardiovascular disease, are associated with vasculature alterations (Beckman & Creager, 2016). The vascular system is particularly sensitive to blood glucose concentrations as endothelial cells (ECs) are in direct contact with the bloodstream and display increased susceptibility due to the dominant expression of insulin-independent glucose transporter GLUT-1(Takata *et al*, 1997), which remains unaffected under high glucose conditions(Kaiser *et al*, 1993).

Hyperglycaemia elicits a collection of molecular alterations in ECs; for instance, glucose-induced activation of NADPH oxidase via PKC induces the production of superoxide radicals and an increase in oxidative stress (Roberts & Porter, 2013). This is further exacerbated by the decreased activity of the antioxidant systems commonly observed in diabetes (Kowluru *et al*, 2007). Additionally, generation of advanced glycation end products (AGEs) promotes a pro-fibrotic state and activation of the RAGE receptor, which activates MAPK and NF-κB pathways (Neumann *et al*, 1999; Ramasamy *et al*, 2008). This leads to the release of pro-inflammatory cytokines, such as IL-6 and TNF-α, and expression of adhesion molecules like ICAM-1 and VCAM-1, ultimately leading to inflammation and vascular damage. Hyperglycaemia-induced ROS also upregulate RAGE (Yao & Brownlee, 2010), and impact the production of nitric oxide (Kolluru *et al*, 2012), leading to vasodilation alterations. Overall, these cumulative effects result in endothelial dysfunction, catalysing the onset of diabetes-associated pathologies in the vasculature.

Longitudinal studies in diabetes patients revealed that past periods of hyperglycaemia can have long-term detrimental effects (Nathan *et al*, 2005; Holman *et al*, 2008), a phenomenon currently known as “metabolic memory”. This concept encompasses the observation that early and rigorous glycaemic control leads to prolonged protection against diabetes complications, even years after discontinuing the intensive glycaemic control (Lachin & Nathan, 2021). A proposed mechanism underlying the metabolic memory posits a positive feedback loop where hyperglycaemia induces overproduction of reactive oxygen species (ROS), which then induce and activate a myriad of pathways that converge in further ROS production (Ihnat *et al*, 2007). This causes continuous activation of the PKC and NF-κB pathways (Giorgi *et al*, 2010; Lingappan, 2018), perpetuating a cycle of ROS-induced cell damage, dysregulation of gene expression and prolonged inflammation even in the absence of hyperglycaemia.

Different studies demonstrated that metabolic memory can be replicated *in vitro.* Transient hyperglycaemia in human cultured cells induce a sustained increase in ROS, overexpression of pro-inflammatory genes and diminished antioxidant response (Zhao *et al*, 2021; Yao *et al*, 2022). Epigenetic alterations are recognized as potential candidates in the establishment and persistence of glucose-induced cellular memory (Siebel *et al*, 2010; Okabe *et al*, 2012; Reddy *et al*, 2015). For example, transient hyperglycaemia in aortic ECs triggers acquisition of active histone marks, while reducing repressive modifications on NOX4 and eNOS promoters, leading to sustained ROS production and vascular dysfunction (Liao *et al*, 2018). Furthermore, NF-κB overexpression is sustained by an antagonistic interplay between histone methyltransferases and demethylases after normalization of glucose, contributing to persistent inflammation (El-Osta *et al*, 2008; Brasacchio *et al*, 2009). Similarly, in renal ECs, a metabolic memory effect has been linked to renal dysfunction both *in vivo* and *in vitro*, linked to reduced chromatin accessibility, alterations in transcription factor binding and diminished histone acetylation (Bansal *et al*, 2020). These alterations also correlate with increased DNA methylation at key genes involved in kidney transport. Moreover, in retinal ECs, hyperglycaemia and ROS production can induce sustained overexpression of DNA (cytosine-5)-methyltransferase 1 (DNMT1), even after normalization of glucose levels (Mishra & Kowluru, 2016), maintaining a pathological epigenetic memory. Additionally, differences in DNA methylation and histone modifications have been observed in pro-inflammatory genes in blood cells of diabetic individuals where a metabolic memory has been established (Miao *et al*, 2014; Chen *et al*, 2016). Nonetheless, the study of the molecular mechanisms driving the establishment and persistence of the metabolic memory is still an active field of research.

Previous studies assessed potential interventions to prevent or “erase” the metabolic memory. For instance, Zhang et al. showed that metformin or resveratrol supplementation in venous ECs prevented the glucose-induced increase in cellular senescence, and that this effect was dependent on increased SIRT1 deacetylation of p53(Zhang *et al*, 2015). More recently, Yao et al. demonstrated that tBHQ-mediated activation of the nuclear factor erythroid 2-related factor 2 (NRF2), a transcription factor that acts as the master regulator of the antioxidant and xenobiotic response in mammals, was able to revert transient high glucose-induced sustained activation of the TGF-beta and NF-kB pathways (Yao *et al*, 2022). In the same study the authors found that treatment with tBHQ reverted collagen accumulation and perivascular fibrosis in a mouse model of metabolic memory. However, the potential beneficial effects of NRF2 activation at a transcriptome-wide scale are yet to be evaluated in the context of the glucose-induced metabolic memory.

Overall, evidence shows that transient periods of pathological hyperglycaemia can imprint a “memory” or “legacy” effect in ECs. This memory encompasses increased inflammation, a surge in oxidative stress, and dysregulation of the genetic programs that ultimately compromise tissue homeostasis. This has important implications in diabetes-related vascular complications and highlights the need for new therapeutic strategies to counteract hyperglycaemia’s legacy effects.

In this study, we characterized the persistent transcriptional and chromatin accessibility alterations induced by transient hyperglycaemia in human endothelial cells. Our findings reveal that the majority of differentially accessible chromatin regions that occur after transient high glucose resulted in a gain of accessibility, with many of these overlapping with putative enhancers that presumably influence the expression of diabetes-related neighbouring genes. Furthermore, our transcription factor motif analysis identified members of the bZIP family as potential players in establishing of the glucose-induced transcriptional legacy response. In accordance, we demonstrated that the pharmacological activation of the NRF2 pathway not only prevented but reverted the glucose-induced transcriptional memory in endothelial cells. Our work contributes to a better understanding of the aetiology of diabetes-associated vascular complications and aids in the development of strategies to mitigate the vascular damage associated with the metabolic memory.

## 2. Results

### 2.1 Transient high glucose exposure induces a metabolic and transcriptional memory in endothelial cells

To explore the persistent effects of transient high glucose concentrations in endothelial cells, we conducted a series of experiments in human umbilical vein endothelial cells (HUVEC) under three treatments: 5.5 mM glucose for 8 days (control), 30 mM glucose for 8 days (HG; high glucose), and 30 mM glucose for 4 days followed by 4 days at 5.5 mM glucose as our memory treatment (Figure 1a). We first evaluated the glycolytic and respiratory rates after transient HG exposure. We found that the extracellular acidification rate (ECAR) – measurement indicative of glycolysis–, was significantly lowered in both our HG and memory treatments, on the other hand, the oxygen consumption rate (OCR) was only decreased in the memory treatment, suggesting that exposure to transient high glucose can induce persistent changes in the basal rates of glycolysis and cellular respiration. (Figure 1b; Supplemental Figure S1a). Notably, we also observed a persistent decrease in the capacity of cells to respond to metabolic stress, as evidenced by lower maximum rates of glycolysis and cellular respiration in the memory treatment compared to control cells (Figure 1b; Supplemental Figure S1a). This suggests that transient hyperglycaemia can establish a metabolic memory under our experimental conditions. To ensure the glucose treatments were not inhibiting cell proliferation, we measured cell growth in our cultures and found that HG exposure had a marginal effect on cell division on days 7 and 8 of culture (Supplemental Figure S1b). Nonetheless, the rate of cell division was maintained at approximately one cell division every 24 hours across all treatments (Supplemental Figure S1c). In line with this, cell cycle assessment showed that only HG treatment had differences in the proportion of cells in each phase compared to the control (Figure 1c): HG-treated cells had an increase of 3.7% of cells in the G2/M phase concomitant with a decrease of 5.1% of cells in the S phase, indicative of a marginal delay in cell cycle. These results indicate that the cells in all our treatments divided approximately 8 times during the course of the 8 days of culture. Additionally, we did not detect differences in cell viability between our treatments as measured by calcein AM assay (Supplemental Figure S1d). All of the above supports the existence of an *in vitro* metabolic memory that is inherited through cell divisions under our experimental conditions. Hyperglycaemia has also been shown to induce persistent oxidative stress, so we measured ROS production in our treatments and found an increase in ROS production in HG, which persisted in the memory treatment (Figure 1d). Consistent with published works (Paneni *et al*, 2012; Yao *et al*, 2022), this finding is indicative that transient hyperglycaemia can induce an oxidative memory in human endothelial cells.

**Figure 1.**
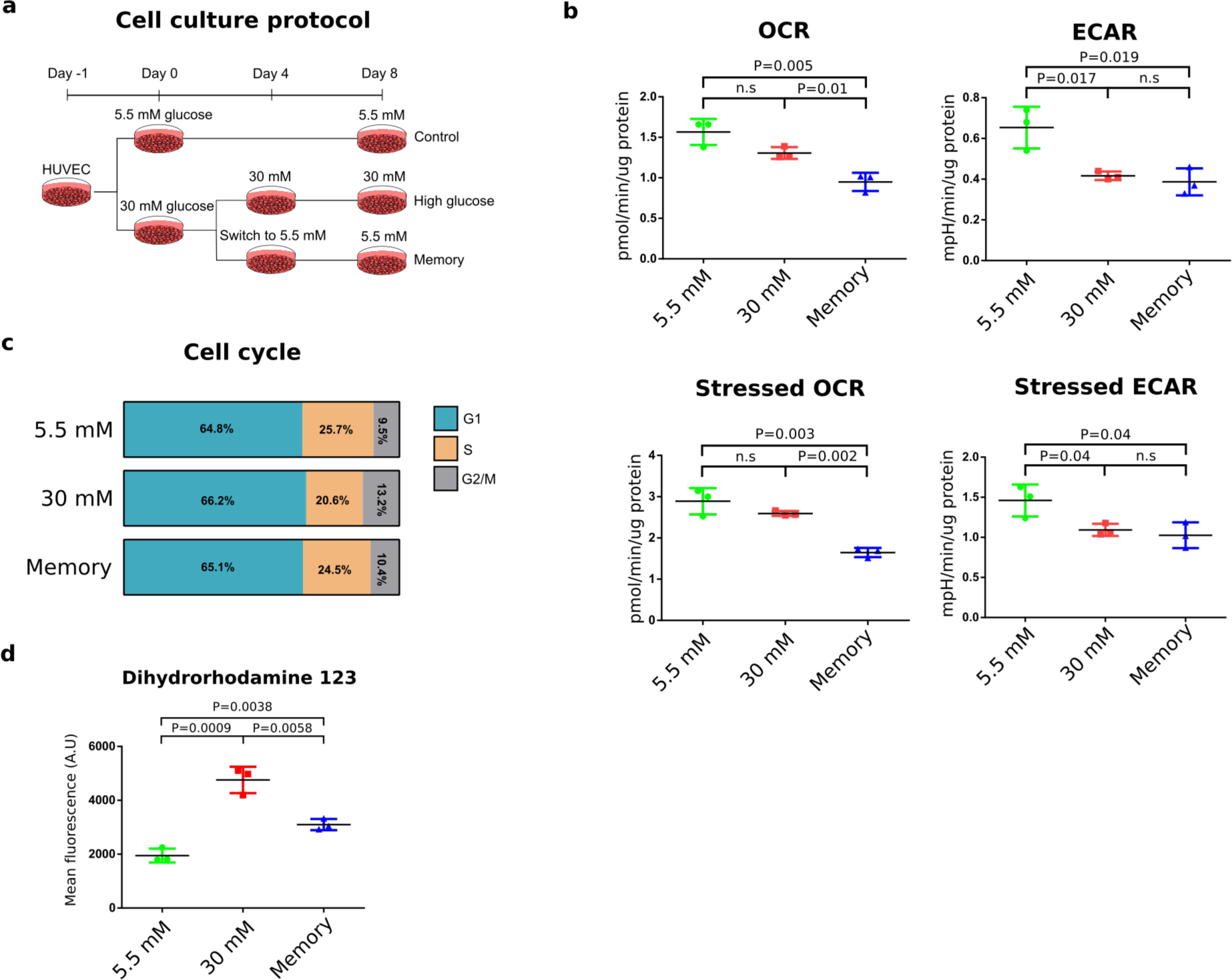
High-glucose-induced metabolic memory in endothelial cells. **(a)** Experimental treatments in HUVECs: Control (5.5 mM glucose, 8 days), high glucose (HG; 30 mM glucose, 8 days), and memory (4 days at 30 mM glucose, then 4 days in 5.5 mM glucose). **(b)** Seahorse quantification of extracellular flux. Basal oxygen consumption rate (OCR) and extracellular acidification rate (ECAR) measurements are followed by stressed measurements made after oligomycin and FCCP injection. Three biological replicates, each with three technical replicates, were used. Differences between pairs of groups were examined with unpaired t-tests. n.s = not significant. **(c)** Percentage of cells in each cell cycle phase evaluated by flow cytometry and DAPI staining. **(d)** Relative ROS concentration measured by dihydrorhodamine 123 via flow cytometry. 3 biological replicates were used per treatment. The principle of the DHR 123 assay is the oxidation of this non-fluorescent reagent by ROS to produce fluorescent rhodamine 123. Differences between pairs of groups were examined by unpaired t-tests.

To further characterize the persistent effects of transient HG in endothelial cells, we analysed the cell transcriptome across treatments. Principal component analysis of the transcriptomic profiles showed that cells exposed to HG and memory treatments were highly similar (Figure 2a; Supplemental Figure S2a). Consistent with this, only 16 differentially expressed genes (DEGs) were found between HG and memory treatments (Supplemental Figure S2b; adjusted p-value <0.05 and log2 fold change > 0.5 and < −0.5; DEGs of pairwise comparisons can be found in the Supplemental Materials). Conversely, in the differential gene expression analysis against the control, we found 1122 DEGs in HG and 1086 DEGs in memory (M) treatment (Figure 2b-c). Of these, 798 (71% of HG-DEGs and 73% of memory-DEGs) were common between HG and memory (henceforth referred to as HG/M-shared DEGs). These results support the existence of a transcriptional memory induced by transient high glucose. We next used the HG-DEGs and memory-DEGs to conduct gene set enrichment analyses to identify overrepresented pathways associated with these experimental conditions. First, we independently examined HG and memory DEGs and found that upregulated HG-DEGs were enriched for terms and pathways known to be affected by hyperglycaemia and diabetes (Yano *et al*, 2004; Coughlan *et al*, 2009; Deshpande *et al*, 2013; Zhang *et al*, 2015; Kyriazis *et al*, 2021), such as the p53, TGF-β, PI3K-AKT, TNF, FoxO and AGE-RAGE pathways, as well as cellular senescence and adhesion molecules, (Figure 2d). Moreover, downregulated HG-DEGs highlighted terms like glutathione metabolism and focal adhesion. On the other hand, upregulated memory-DEGs were enriched for terms such as cellular senescence, and p53 and TGF-beta pathways, while downregulated memory-DEGs showed terms such as focal adhesion and the PI3K-Akt and NRF2 signalling pathways (Figure 2d). Many of these pathways were confirmed using the GSEA (Subramanian *et al*, 2005) software analysis (Supplemental Figure S2c).

**Figure 2.**
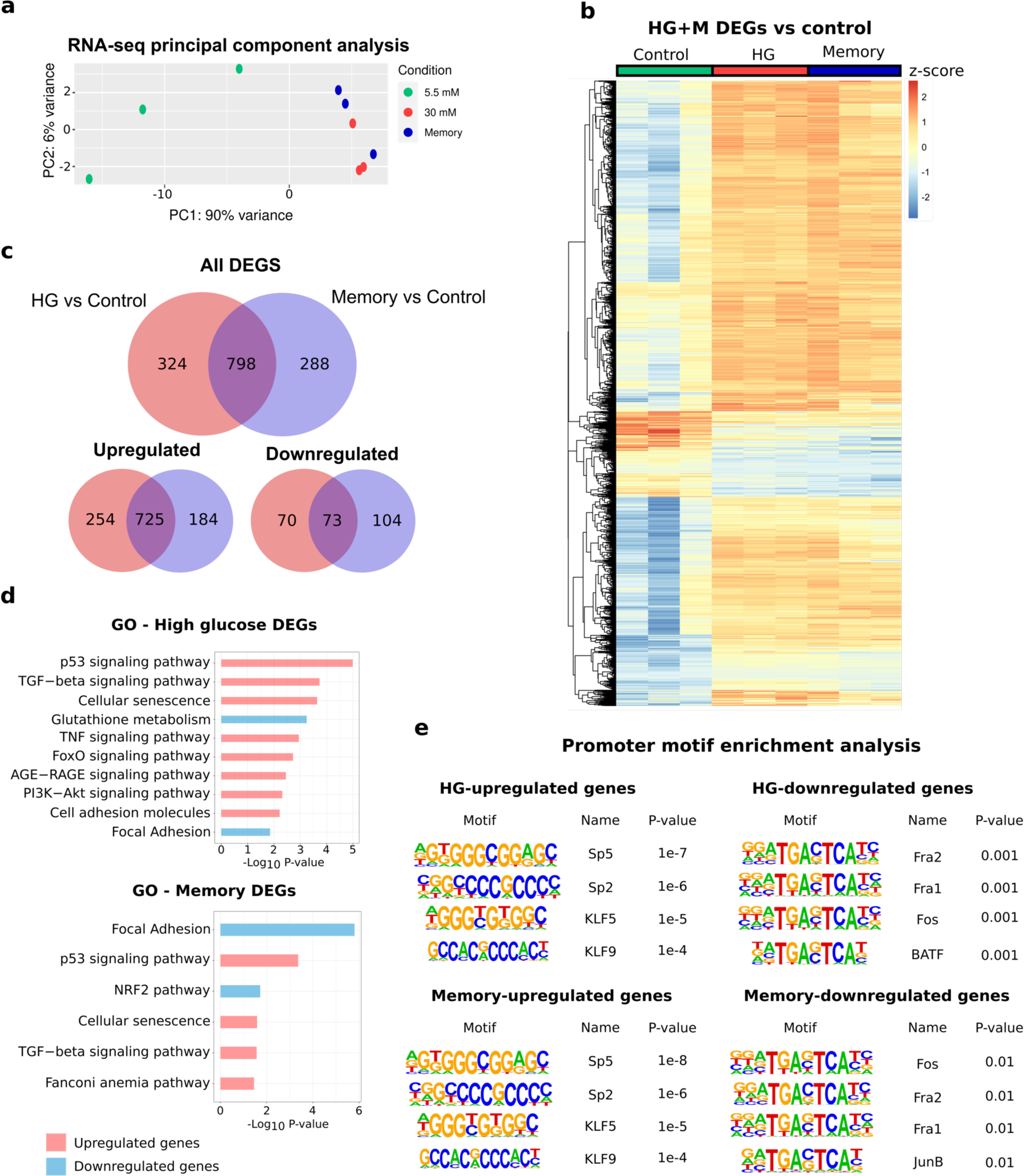
High-glucose-induced transcriptional memory in endothelial cells. **(a)** Principal component analysis of RNA-seq samples with three biological replicates per treatment. **(b)** Heatmap displaying the relative expression changes of 1410 differentially expressed genes (DEGs): HG vs control + memory vs control. **(c)** Venn diagram showing the overlap between HG vs. control and memory vs control DEGs. **(d)** Pathway enrichment analysis of HG vs. control and memory vs. control DEGs. **(e)** Transcription factor motif enrichment analysis conducted in promoters of HG vs. control and memory vs control DEGs. A window ranging from −1000 to +100 bp from the TSS was used for the search.

We next examined the 798 HG/M-shared DEGs, and we found that upregulated HG/M-shared DEGs were enriched for terms such as p53 and TGF-beta pathways (Supplemental Figure S3a). Furthermore, downregulated HG/M-shared DEGs were enriched for terms of focal adhesion and PI3K-AKT signalling pathway. To evaluate the distinctive transcriptomic profile of transient hyperglycaemia exposure, we examined independently both the 324 DEGs unique to HG and the 288 DEGs exclusive to the memory treatment (“HG-unique” and “memory-unique”, respectively). Interestingly, upregulated HG-unique genes were enriched for terms like the TNF, TGF-β and AGE-RAGE pathway as well as cellular senescence and adhesion molecules (Supplemental Figure S3c). In contrast, downregulated HG-unique genes were enriched for glutathione metabolism. On the other hand, upregulated memory-unique genes were enriched for the Fanconi anaemia pathway, linked to DNA repair and replication, whereas memory-unique downregulated genes were enriched for focal adhesion and the NRF2 pathway (Supplemental Figure S3c).

To gain insights into the transcriptional regulatory program underlying metabolic memory, we conducted analyses of possible transcription factors (TFs) driving persistent transcriptional changes. To this end, we examined the promoter sequence of HG and memory DEGs to perform a TF motif enrichment query. Our analysis revealed that the KLF/SP family of transcription factors exhibited the most statistically enriched motifs in the upregulated genes of both HG and memory (Figure 2e). This finding aligns with our differential expression analysis, which identified KLF3, KLF5, KLF9, and SP4 as persistently upregulated in the memory treatment (Supplemental Figure S4c). In contrast, the promoters of downregulated HG and memory genes were enriched for the predicted motifs of FRA1, FOS and JUN (Figure 2e). Of note, FRA1 (*FOSL1*) was downregulated in HG and memory treatments (Supplemental Figure S4c). We noticed that these motifs share the same core sequence “TGA(C/G)TCA” which corresponds to the common binding motif of the basic leucine zipper (bZIP) domain transcription factors (Rodríguez-Martínez *et al*, 2017). This family includes the nuclear factor erythroid 2-related factor 2 (NRF2) transcription factor, known to be the master regulator of the antioxidant response in mammals. Collectively, these findings suggest that transient high-glucose triggers a persistent transcriptional memory in human endothelial cells, with bZIP transcription factors potentially playing an important role.

### 2.2 Transient high-glucose treatment induces persistent chromatin accessibility changes in non-promoter regions

To further investigate the mechanisms underlying glucose-induced transcriptional memory, we performed ATAC-seq to assess potential chromatin accessibility changes associated with the HG and memory treatments. We identified 79,376 unique peaks, with 45,363 shared across treatments (Supplemental Figure S5a). The distribution of called peaks was consistent across conditions, with the majority located in introns, followed by intergenic and promoter regions (Supplemental Figure S5b). Analysis of the ATAC-seq signal around peaks by genomic feature revealed an increase in accessibility in intergenic and intronic regions in HG and memory, which was absent in peaks at promoters or exons (Supplemental Figure S5d). Differential chromatin accessibility analysis of HG versus control identified 2,863 differentially accessible regions (DARs; fold change >1.5 and p-value <0.0001); over 90% of which were located in either intronic or intergenic regions (Figure 3a). Among these DARs, 2,641 (92%) had increased accessibility in HG versus control (Figure 3c). Similarly, in the memory versus control comparison, 1869 DARs were identified with 89% of them located in either intronic or intergenic regions (Figure 3a). Additionally, the majority of memory-DARs (81%) had increased accessibility compared with the control (Figure 3c). Notably, inspection of the mean ATAC signal across the 4166 HG plus memory DARs revealed that changes in these regions were not fully reverted to control levels in the memory treatment (Figure 3c), indicating the establishment of an epigenetic memory induced by transient hyperglycaemia.

**Figure 3.**
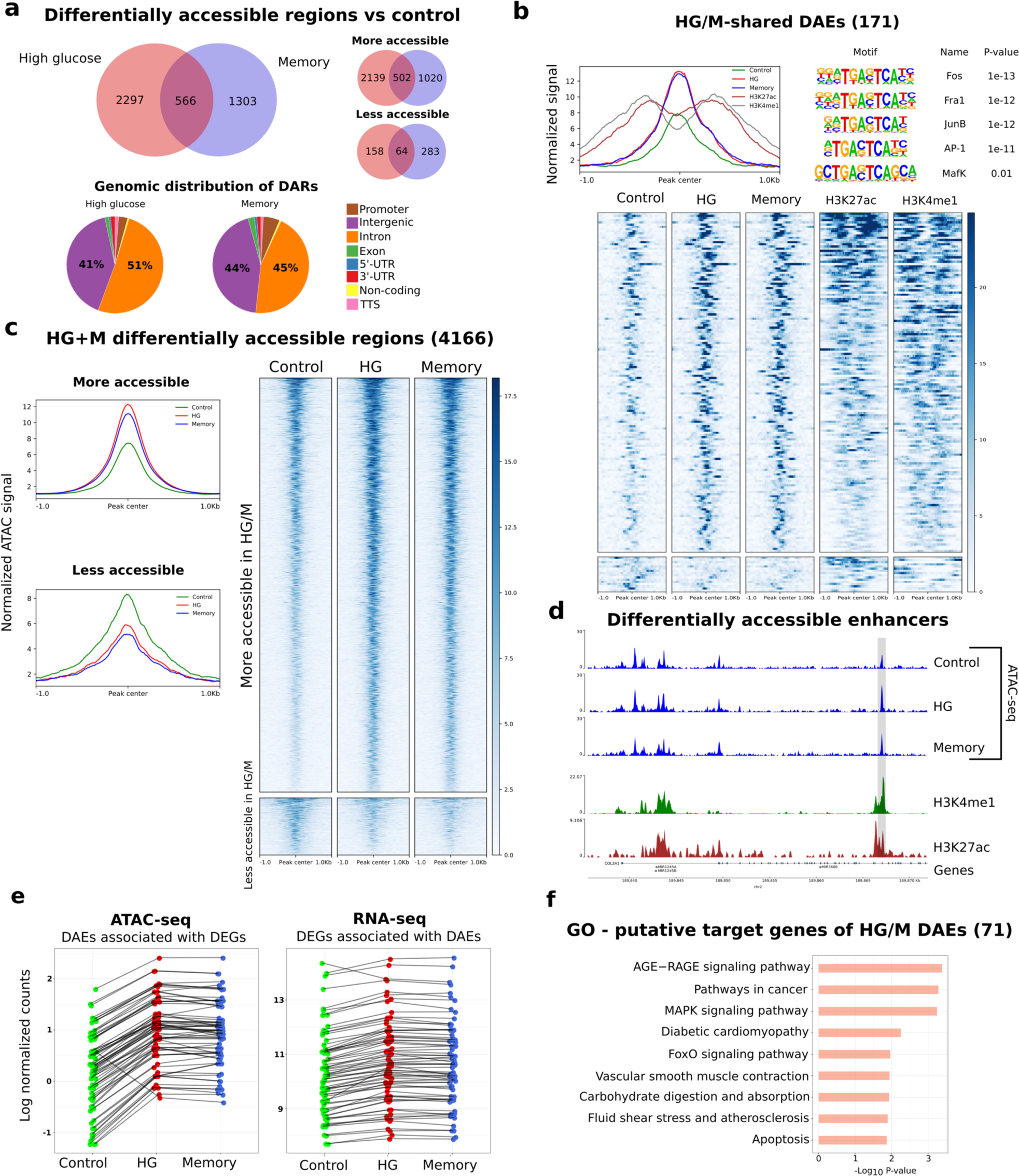
Transient high glucose induces persistent changes in HUVEC chromatin accessibility. **(a)** Top: Venn diagram showing the overlap between high glucose (HG) vs. control and memory (M) vs. control Differentially Accessible Regions (DARs). Bottom: pie chart showing the genomic distribution of DARs. **(b)** Left: ATAC-seq signal plot showing the control, HG and memory, alongside H3K4me1 and H3K27ac ChIP-seq signal in Differentially Accessible Enhncers (DAEs). Right: TF motif enrichment analysis in DAEs within ±50 bp from the ATAC-seq peak center. Bottom: heatmap of DAEs showing the control, HG and memory ATAC-seq signal along with the H3K4me1 and H3K27ac ChIP-seq signal. **(c)** Left: normalized ATAC-seq signal by genomic feature in control, HG and memory samples. Right: heatmap of ATAC-seq signal in HG+M DARs in control, HG and memory samples. **(d)** Example of a differentially accessible enhancer located in an intron of the COL3A1 gene highlighted in gray; note that chromatin accessibility is not reverted to the control levels in the memory treatment. **(e)** Association between changes in chromatin accessibility in DAEs and changes in gene expression of putative target DEGs identified by the GREAT software. **(f)** Pathway enrichment analysis of the 71 HG/M DEGs that are putative targets of DAEs.

Given that the majority of the identified DARs were in intergenic and intronic regions and considering that genomic long-range interactions play critical roles in fine-tuning gene expression in response to the environment (Maurya, 2021), we explored if non-promoter HG/M-shared DARs could be annotated to enhancer regions. Using HUVEC ChIP-seq data from ENCODE for two enhancer-enriched histone marks, H3K4me1 and H3K27ac, we found that 171 out of the 566 HG/M-shared DARs were enriched for both histone marks, so we catalogued them as differentially accessible putative enhancers (DAEs; Figure 3b, d). Next, we employed the Genomic Regions Enrichment of Annotations Tool (McLean *et al*, 2010) to identify potential target genes of DAEs, finding that 71 out of the 297 (24%) putative target genes were also identified as DEGs in our RNA-seq (Supplemental Figure S5c). Remarkably, the direction of transcriptional change corresponded with the DAEs accessibility trend (Figure 3e). Pathway analysis of these putative target genes highlighted terms such as the AGE-RAGE, MAPK and FoxO signalling pathways (Figure 3f). Finally, to identify candidate TFs driving the glucose-induced chromatin accessibility changes, we searched for enriched motifs within DAEs. We found that DAEs were significantly enriched for motifs of bZIP transcription factors, including FOS, FRA1, JUN and MAFK (Figure 3b). To note, MAFK is a partner of NRF2 that participates in cellular oxidative stress response (Blank, 2008). These findings are consistent with our earlier motif enrichment analysis of downregulated HG and memory genes that suggested that bZIP transcription factors influence the memory transcriptional response.

In summary, we found that transient hyperglycaemia induces persistent chromatin accessibility changes mainly in non-promoter regions, including putative enhancers, which could contribute to the glucose-induced transcriptional changes. Furthermore, bZIP transcription factors emerge as potential regulators of these processes.

### 2.3 Activation of NRF2 pathway reverts the glucose-induced transcriptional memory

Our findings indicated that HG and memory treatments presented downregulation of antioxidant response-related genes, including those in the glutathione metabolism and NRF2 pathway. Additionally, we observed bZIP motif enrichment at promoters of downregulated memory-DEGs and HG/M-shared differentially accessible enhancers. These prompted us to explore the role of the transcription factor NRF2 in the persistent effects of transient high-glucose in endothelial cells. We first used lentiviral vectors to express GFP (as control) or human NRF2 in HUVECs, and then exposed these cells to the HG and memory treatments to evaluate transcriptional changes via RNA-seq (Figure 4a). To confirm NFR2 overexpression (OE), we validated its expression in our transduced RNA-seq samples, finding a >10-fold increase in transcript abundance compared with the control (Supplemental Figure S4c). Then, we evaluated the effect of NRF2 OE during HG and memory treatments by analysing changes in expression of their corresponding subsets of DEGs. We found that only 26% (281 of 1086) of the memory-DEGs, and 20% (220 of 1122) of the HG-DEGs remained differentially expressed compared to the control after NRF2 OE (Figure 4b-c; Supplemental Figure S4d). These results indicate that NRF2 OE effectively mitigates the transcriptional effects of HG exposure. Subsequently, we performed pathway enrichment analysis on both NRF2 OE-rescued and non-rescued genes. Interestingly, NRF2 OE-rescued genes were enriched for terms related to cellular senescence, Fanconi anaemia, focal adhesion, and glutathione metabolism, as well as the p53 and FoxO pathways (Supplemental Figure S3d). Meanwhile, NRF2 OE-not rescued genes showed terms related to cellular senescence, adhesion molecules and the TGF-beta, p53, and NF-κB pathways, indicative of NRF2 OE being incapable of reverting changes in pro-inflammatory genes.

**Figure 4.**
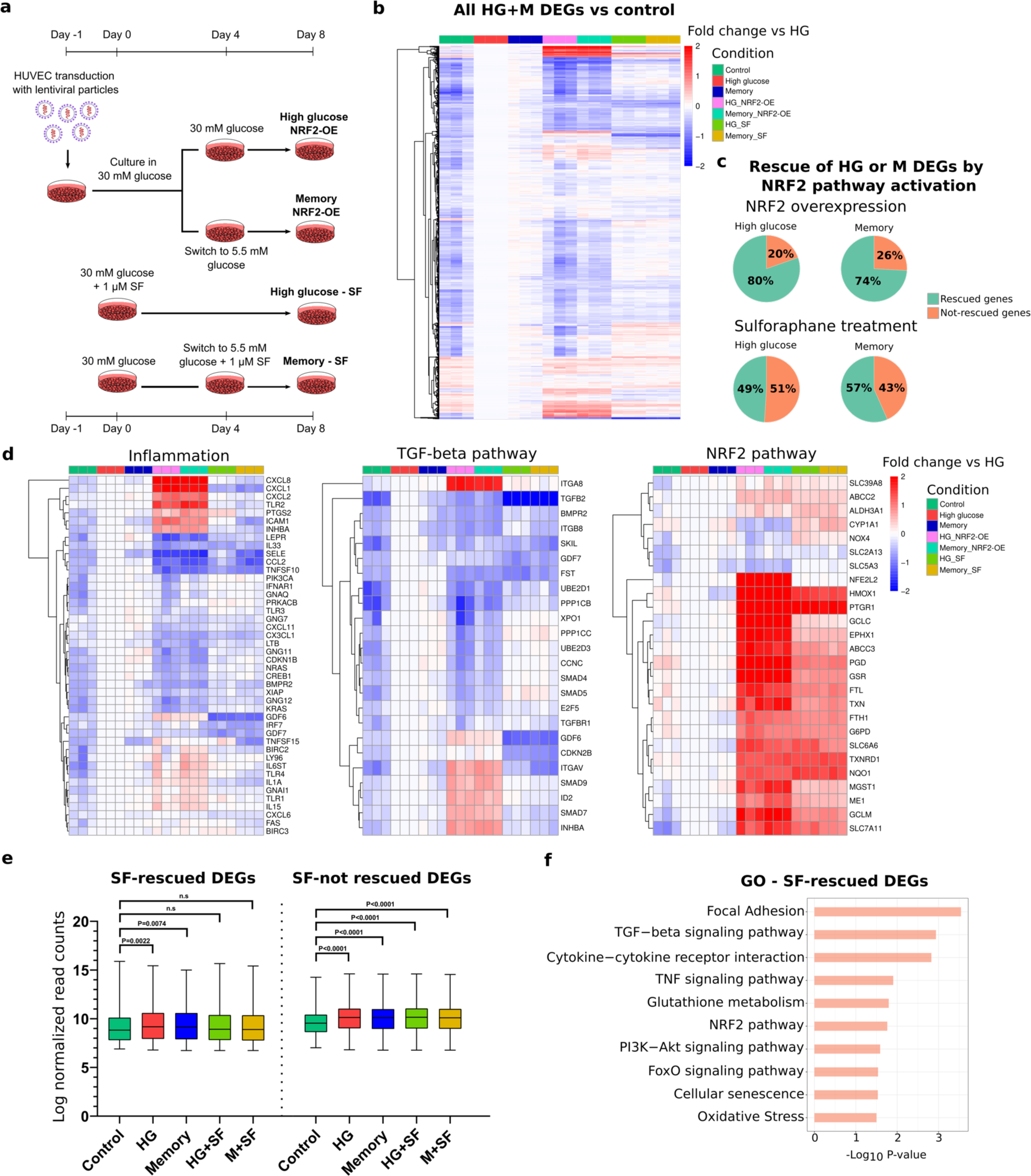
NRF2 pathway activation via gene overexpression or sulforaphane supplementation reverts high glucose-induced transcriptional changes. **(a)** Protocol used for NRF2 overexpression (OE) or sulforaphane (SF) supplementation in HUVEC. For NRF2 OE, cells were transduced with NRF2-carrying lentiviral particles (MOI=20) while cultured in 5.5 mM glucose. After 24 hours, glucose concentration was increased to 30 mM for 8 days (HG treatment), or 4 days in 30 mM followed by 4 days in 5.5 mM of glucose for memory treatment. For SF treatments, 1 μM SF was added during the 8-day HG treatment or at the switch time point of the memory treatment (final 4 days in 5.5 mM glucose). This allowed us to test if SF could prevent and/or revert the glucose-induced transcriptional memory. **(b)** Heatmap depicting the effect of NRF2 activation by OE or SF treatment on HG+M DEGs. Data is shown as fold change versus HG. **(c)** Percentage of DEGs found in HG or memory treatments that are rescued via NRF2 pathway activation by NRF2 OE or SF supplementation. **(d)** Heatmaps of HG/M DEGs grouped in three categories: inflammation, TGF-beta pathway and NRF2 pathway. Data is shown as fold change versus HG. **(e)** Plot showing the changes in expression of HG/M DEGs after SF treatment, distinguishing between those rescued and not rescued by SF. Differences between groups were examined by unpaired t-tests. n.s = not significant. **(f)** Pathway enrichment analysis of HG/M DEGs rescued by sulforaphane supplementation.

To circumvent the disadvantages of lentiviral systems for therapeutic purposes, we used sulforaphane (SF) to induce the NRF2 pathway. SF exerts NRF2 activation by inhibiting its interaction with the repressor KEAP1, leading to NRF2 accumulation and nuclear translocation^37^. We tested various SF concentrations and determined that 1 μM effectively induced the expression of NRF2 target genes HMOX1 and NQO1 without impacting cell viability or proliferation (Supplemental Figure S1b-d; Supplemental Figure S4a). In agreement with the mechanism of NRF2 activation by SF, culture of HUVEC with 1 μM SF did not induce NRF2 transcription changes (Supplemental Figure S4a). Moreover, RT-qPCR of selected HG/M DEGs confirmed that 1 μM SF reverted their expression changes in our HG and memory treatments (Supplemental Figure S4b), hence we used this concentration for subsequent experiments.

We performed RNA-seq on cells exposed to our HG and memory treatments, adding SF throughout the entire HG treatment, or after switching to normal glucose in the case of the memory treatment (Figure 4a). This allowed us to evaluate if SF could prevent or reverse the glucose-induced transcriptional memory. Remarkably, SF ameliorated the transcriptional alterations in both the HG and memory treatments, where only 471 out of 1086 (43%) memory-DEGs and 572 out of 1122 (51%) HG-DEGs remained differentially expressed after SF treatment (Figure 4b-c, e). Interestingly, upon examining the SF-rescued genes we found notable differences in the enriched pathways compared with those of NRF2 OE-rescued genes: SF-rescued genes highlighted terms such as TGF-beta pathway, cellular senescence, and cell adhesion molecules, which were absent in the NRF2 OE-rescued genes (Figure 4f). Additional terms enriched in the SF-rescued genes included the p53, PI3K-Akt, TNF, AGE-RAGE, and NRF2 pathways, as well as focal adhesion and glutathione metabolism. On the other hand, genes not rescued by SF supplementation showed terms like cellular senescence and the p53 and MAPK pathways (Supplemental Figure S3b). The effect of NRF2 activation, either by NRF2 OE or SF treatment, on the expression of DEGs belonging to selected pathways is shown in Figure 4d.

Taken together, our results show that NRF2 activation effectively prevents most transcriptional alterations induced by both persistent and transient high glucose exposure. In addition, NRF2 activation by sulforaphane can either prevent or revert the glucose-induced transcriptional memory in endothelial cells, avoiding the disadvantages of lentiviral vectors and risks of overexpressing a transcription factor beyond the physiological range. These findings suggest that modulating the NRF2 pathway could be a promising therapeutic strategy to erase the metabolic memory induced by hyperglycaemia.

## 3. Discussion

Transient pathological hyperglycaemia can alter vascular homeostasis by inducing sustained changes in gene expression, such as upregulation of pro-inflammatory pathways and increased oxidative stress (Brasacchio *et al*, 2009; Paneni *et al*, 2012). Here, we report that activation of the NRF2 pathway, either via genetic overexpression or pharmacological activation with sulforaphane, reverts the high-glucose-induced transcriptional memory in human endothelial cells. Remarkably, sulforaphane not only prevents most of the prolonged transcriptional alterations caused by HG, but it can also revert them once established. Sulforaphane is a plant-derived isothiocyanate found in cruciferous vegetables, possessing a diverse array of biological effects (Mangla *et al*, 2021). For instance, sulforaphane blocks inflammation by inhibiting NF-kB activity, and has antioxidant properties by modifying cysteine residues of KEAP1, the inhibitor of NRF2, resulting in NRF2 stabilization and nuclear translocation, where it binds to the antioxidant response element (ARE) motif sequence to activate antioxidant and stress response genes(Hu *et al*, 2011). While NRF2-independent chemoprotective mechanisms for sulforaphane have been described (Greaney *et al*, 2016), our data showing that NRF2 gene overexpression resulted in a similar outcome suggests that the beneficial effects conferred by sulforaphane in our HG and memory treatments occur mainly through the activation of NRF2 pathway.

Interestingly, while NRF2 OE restored the expression of more DEGs in either HG or memory compared to sulforaphane treatment, our RNA-seq analysis revealed that NRF2 OE led to increased expression of pro-inflammatory genes, surpassing even the levels observed in HG/M alone in some cases (see “inflammation” in Figure 4d). We hypothesize that this effect resulted from lentiviral-mediated overexpression of NRF2 beyond physiological range, an effect not observed with sulforaphane. In line with this, previous studies reported that NRF2 can activate pro-inflammatory genes in some contexts (Wruck *et al*, 2011). In fact, NRF2 ablation is reported to decrease NF-kB subunits p50 and p65 in mouse fibroblasts (Yang *et al*, 2005), highlighting the importance of TF abundance balance in modulating its function and binding to target genes.

Many unresolved questions about the metabolic memory revolve around the molecular basis driving the establishment and persistence of glucose-induced transcriptional memory. Epigenetic regulation of gene expression stands out as a possible mechanism for the modulation of these processes (Reddy *et al*, 2015; Dhawan *et al*, 2022). With that in mind, we assessed the chromatin accessibility landscape of HUVEC exposed to our HG and memory treatments, finding that hyperglycaemia led to changes in accessibility specifically within intergenic and intronic regions that are not reversed post-glucose normalization. Interestingly, most of these DARs resulted from an increase in accessibility, and when we overlapped them with HUVEC’s H3K4me1 and H3K27ac ChIP-seq data, we found that a proportion of those regions were *bona fide* enhancers. Moreover, we discovered that about 25% of the putative target genes of those DAEs were HG/M-DEGs, suggesting that enhancer dynamics contribute to the glucose-induced transcriptional alterations. Our observation that enhancers retain an accessibility memory is relevant, given their role in controlling the magnitude and timing of gene expression (Maurya, 2021). Based on their epigenetic signatures, enhancers can reside in “active”, “inactive” or “poised” states (Creyghton *et al*, 2010). Thus, we hypothesize that transient hyperglycaemia impacts the epigenetic and functional state of enhancers, priming them to amplify or sustain the transcriptional changes. This mechanism mirrors how inflammation can imprint an enhancer’s epigenetic memory in immune and endothelial cells (Drummer *et al*, 2021). Ergo, in diabetes patients, repetitive cycles of pathological hyperglycaemia could set enhancers into a pathological-memory state.

Regarding glucose-induced transcriptional changes, we found a differential promoter enrichment of TF motifs between our HG/M upregulated and downregulated genes. Specifically, upregulated genes were enriched in motifs of the KLF/SP family of TFs, while downregulated genes favoured members of the bZIP family. Concordantly, we observed upregulation of KLF3, KLF5, KLF9 and SP4 in HG and memory treatments, and notably, all of these genes were rescued by sulforaphane treatment, suggesting that one possible mechanism by which NRF2 reverts the glucose-induced transcriptional memory is through the restoration of the expression of these transcription factors. Members of the KLF/SP family are known to be involved in cell differentiation, cell proliferation and metabolism. Interestingly, elevated KLF5 expression has been linked to oxidative stress, vascular remodelling and impaired lipid metabolism (Kyriazis *et al*, 2021).

On the other hand, promoters of downregulated HG/M-DEGs exhibited enrichment for bZIP family members, which share common features in their binding motifs and genomic targets (Rodríguez-Martínez *et al*, 2017). Notably, we identified bZIP member FRA1 consistently downregulated in our HG and memory treatments, and similarly to KLF/SP TFs, FRA1 expression was restored after sulforaphane treatment. FRA1 is a member of the AP-1 complex known to induce the expression of antioxidant genes such as HMOX1 and sulfiredoxin (Lee *et al*, 2000; Soriano *et al*, 2009) —genes persistently downregulated in our HG and memory treatments and modulated by NRF2. This observation, together with previous reports showing that NRF2 protein levels are persistently diminished after transient HG(Yao *et al*, 2022), could explain the paradoxical finding in various studies(Hodgkinson *et al*, 2003; Kowluru *et al*, 2007; Sekhar *et al*, 2010), including our own, regarding decreased expression of antioxidant genes under HG. Nonetheless, research on the interplay between these TFs regarding their activities and shared molecular targets is still lacking.

Hyperglycaemia can disrupt glucose metabolism by inhibiting glycolysis and pentose phosphate pathways (Du *et al*, 2000; Zhang *et al*, 2000), while increasing ROS by reduced NADPH levels, accumulation of glycolysis intermediates and subsequent augmented flux of alternative metabolic routes like the polyol and hexosamine pathways (Giacco & Brownlee, 2010). Concordantly, we observed that transient HG persistently reduced glycolysis and oxygen consumption. Similar results were observed in retinal ECs where prolonged HG resulted in decreased glycolysis (Bertelli *et al*, 2022), while in HUVEC, transient HG induced a persistent decrease in oxygen consumption(Yao *et al*, 2022), a result recapitulated in our data. Additionally, HUVEC exposed to AGEs also exhibited reduction in glycolysis and cellular respiration (Li *et al*, 2017). Notably, our RNA-seq data revealed increased expression of pentose phosphate pathway genes such as G6PD (glucose-6-phosphate dehydrogenase), PGD (6-phosphogluconate dehydrogenase) and TALDO1 (transaldolase 1) following sulforaphane treatment, which may contribute to the beneficial effects we observe with the supplementation after HG and memory treatments.

Overall, our work demonstrates that transient hyperglycaemia induces prolonged alterations in the transcriptome, oxidative stress, glucose metabolism, and chromatin accessibility in endothelial cells, but some limitations should be considered. First, while we exposed cells to normal glucose after an equivalent time in HG (4 days), we did not evaluate extended recovery times and the duration of the memory effect. Other authors have shown that the metabolic memory can persist for up to 11 days *in vitro* (Roy et al, 1990). Thus, it would be important to characterize the potential weakening of the memory. Second, in our study, endothelial cells underwent four divisions after the hyperglycemic stimulus, yet cell division rate in adults is slower and influenced by age (Hobson & Denekamp, 1984; Hoshi & McKeehan, 1986). Hence, it is difficult to extrapolate the time required to establish a glucose-induced metabolic memory in the adult vasculature. Additionally, while our study focused on constant hyperglycaemia, patients with diabetes experience drastic glucose concentration fluctuations, which have been associated with poor outcomes (Zhou *et al*, 2020). Future research investigating the impact of these pathological fluctuations in the establishment of the cell memory would be important. Third, our results revealed that sulforaphane could reverse the transcriptional changes associated with the glucose-induced memory. Still, it would be relevant to assess - especially *in vivo* - whether this reversal is accompanied by whole-genome epigenetic changes, in particular, histone modifications and 3D chromatin architecture. Lastly, while we verified that NRF2 pathway activation can prevent and revert the transcriptional changes caused by transient HG, our data are insufficient to determine if this effect results from direct action of this TF and/or from downstream regulation of NRF2 targets.

The metabolic memory phenomenon has been studied for over three decades (Lachin & Nathan, 2021), yet currently there are no specific treatments to ameliorate the diabetes-associated vascular complications, which comprise the leading causes of morbidity and mortality in patients with this disease. Our study highlights the potential use of sulforaphane to revert the glucose-induced transcriptional memory in human endothelial cells, and while other compounds have antioxidant and anti-inflammatory properties, sulforaphane is among the ones with the highest bioavailability(Houghton *et al*, 2016) and offers additional health benefits, including anti-cancer and anti-inflammatory properties (Mangla *et al*, 2021). In addition, small-scale studies using sulforaphane in diabetes patients have shown promising results (Axelsson *et al*, 2017).We look forward to future randomized longitudinal studies with larger cohorts that will assess the efficacy and safety of long-term sulforaphane supplementation in mitigating the impact of diabetes-associated metabolic memory.

Taken together, our results demonstrate that transient hyperglycaemia induces sustained transcriptional, metabolic and epigenetic alterations in human endothelial cells. Our study also highlights the potential use of sulforaphane to revert the vasculature transcriptional memory in the context of diabetes.

## 4. Materials and methods

### 4.1 Cell culture

Pooled-donor human umbilical vein endothelial cells (HUVEC) (Lonza, Basel, Switzerland; C2519A) were cultured in endothelial basal medium EBM (Lonza, Basel, Switzerland; CC-3121) supplemented with SingleQuots (Lonza, Basel, Switzerland; CC-4133) at 37°C, 5% CO_2_, and 8% O_2_ for no more than 8 passages, using 0.05% trypsin to subculture the cells. HUVEC were seeded in pre-coated gelatine plates; medium was replaced every 48h. Cells were cultured for 8 days in glucose 5.5 mM (control), 30 mM (high glucose), or memory treatment (4 days at 30 mM, then 4 days at 5.5 mM). Seeding number was adjusted to reach 70-80% confluency after 4 days of culture, and only one passage was performed throughout the 8 days of treatment (at day 4). 1 μM sulforaphane (Sigma, S4441) was added into media from a 2.8 mM stock. Supplementation was maintained for 8 days in 5.5- or 30-mM glucose, or when cells switched to 5.5 mM in the memory treatment. Controls received equivalent volume of vehicle (DMSO).

### 4.2 Extracellular flux assays

Extracellular acidification rate (ECAR) and oxygen consumption rate (OCR) were measured using the Seahorse XFe96 analyser (Agilent, Santa Clara, CA) with Cell Energy Phenotype Test kit. The day before, 10,000 HUVEC/well from control, high glucose or memory treatments were seeded in gelatine pre-coated 96-well plate and cultured in EBM with the corresponding glucose concentration at 37°C, 5% CO_2_, and 8% O_2_. On assay day, cells were washed with 200 μL assay medium (Seahorse XF Base Medium with 1 mM pyruvate, 2 mM glutamine, and 10 mM glucose) and incubated in 180 μL of fresh assay medium for 1 h at 37°C before the assay. The Cell Energy Phenotype test consists of three baseline measurements of oxygen consumption rate (OCR) and extracellular acidification rate (ECAR) followed by five stressed measurements. The later are induced by the simultaneous addition of oligomycin, a complex V inhibitor, and carbonyl cyanide-p-trifluoromethoxyphenylhydrazone (FCCP), an uncoupling agent. Oligomycin and FCCP were added to each well to a final concentration of 1 mM and 0.5 mM, respectively. Results were analyzed with Agilent’s Wave software and normalized to protein concentration. Three biological, and three technical replicates were used per condition.

### 4.3 Cell viability assays

For Calcein AM (ThermoFisher Scientific, Waltham, MA; C3100MP), cells were harvested after treatments and washed with PBS, then resuspended in HBSS with 1 μM Calcein AM for 20 min at 37°C protected from light. Dye was then removed and fresh HBSS was added. Fluorescence intensity was measured with FACSMelody (BD, Franklin Lakes, NJ). Single cells were gated, and 10,000 cells were acquired per sample to identify high-fluorescent viable cells. Unstained cells were used as controls. Data were analysed with FlowJo. Three biological replicates were acquired from each treatment.

### 4.4 Cell growth curves and cell cycle

HUVEC were seeded on gelatine coated 12-well plates and counted daily for 8 days using the CytoSmart counter (Corning, Corning, NY). Three biological replicates with 2 technical replicates each were evaluated per day per treatment. At day 4, cells were subcultured. To maintain a consistent curve, cell counts from days 5-8 were calculated using a division rate derived from day 4 cell counts. This rate was the day’s total cells divided by starting cells from day 4 subculture.

To obtain the overall mean division rate, we used the following formula:

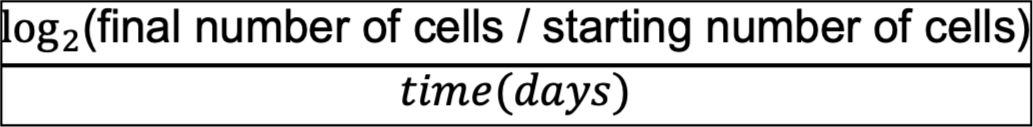

For cell cycle analysis, cells were trypsinized and resuspended in DPBS with 0.1% NP-40, and DAPI (1 ug/mL). After 10 minutes incubation, cell DNA content was measured with FACSMelody (BD) using UV laser on a linear scale. 5,000-10,000 single cells were acquired per sample with three biological replicates per treatment. FlowJo software was used to calculate the proportion of cells in each cell cycle phase.

### 4.5 ROS assay

Reactive oxygen species were determined using Dihydrorhodamine123 (DHR 123; ThermoFisher Scientific). After 8 days, cells were trypsinized, washed with PBS, and resuspended in HBSS with 1 μg/mL DHR 123. After 20 min at room temperature, cells were rinsed again with DPBS and resuspended in HBSS. Fluorescence was measured with Attune NxT flow cytometer (ThermoFisher Scientific). 5,000-10,000 single cells were acquired per sample with three biological replicates per treatment. Unstained cells served as controls. Analysis was done using FlowJo software, with mean fluorescence indicating ROS levels.

### 4.6 RNA-seq library preparation

HUVEC total RNA was isolated using the QuickRNA Microprep kit (Zymo Research, Irvine, CA) following manufacturer’s instructions. RNA’s integrity and concentration were evaluated on a 2100 Bioanalyzer (Agilent). Samples with a RIN >9 were selected for RNA-seq library preparation. Libraries were prepared from 1 μg of RNA using the TruSeq Stranded mRNA library prep kit (Illumina) following the reference guide. Synthesized libraries were sequenced on a Nextseq 500 System (Illumina) with paired-end 75 bp reads. Summary of sequenced and mapped reads are in Table 1.

### 4.7 ATAC-seq library preparation

Libraries for Assay for transposase-accessible chromatin were prepared as described previously (Buenrostro *et al*, 2015). Briefly, 50,000 HUVEC were harvested, resuspended in ice-cold DPBS, and centrifuged (1 min at 1300 x g, 4°C). Pelleted cells were resuspended in 50 μL of cold lysis buffer and centrifuged (10 min at 1200 x g, 4°C). After discarding supernatant, 50 μL of transposase mix with 2.5 μL of Tn5 transposase (Illumina, San Diego, CA) were added and incubated for 30 min at 37°C. The reaction was stopped by purification with MinElute PCR purification kit (Qiagen, Hilden, Germany) following manufacturer’s instructions. Library amplification by PCR was done after determining number of cycles needed. Products were size-selected with Agencourt AMPureXP beads (Beckman Coulter, Brea, CA) and sequenced on a Nextseq 500 System (Illumina, 75 bp paired-end reads). Details are shown in Table 2.

### 4.8 RNA-seq data analysis

Data quality was assessed using FastQC. Alignments of paired-end samples were generated using STAR v2.7.3a with default parameters using the human hg38 genome and annotations from the UCSC portal. Raw gene count matrices were generated using FeatureCounts with flags “-p -s 2”. Differential gene expression was analysed with DESeq2 v1.36; genes with an adjusted p-value <0.05 and a log2 fold change >0.5 and <-0.5, were considered differentially expressed. Pathway enrichment was done via the EnrichR web tool with a 0.05 p-value cut-off to consider a term to be enriched. The Kyoto Encyclopedia of Genes and Genomes (KEGG) and the Wikipathways databases were used for our analyses. Motif enrichment analysis in promoters of DEGs was done with the findMotifs tool from HOMER software with flags “-start −1000 -end 100”. Heatmaps were generated using DESeq2 variance stabilizing transformation normalized counts and the R package Pheatmap. Volcano plots were generated using the R package EnhancedVolcano.

### 4.9 ATAC-seq data analysis

Data quality was assessed with FastQC. Adapters were trimmed using TrimGalore, and alignment files were referenced to the human hg38 genome with Bowtie2 v2.3.5 with flag “-X 1000” to align fragments up to 1 kb in length. Mitochondrial and PCR duplicate reads were discarded using Samtools and PicardTools, respectively. ENCODE blacklist-mapped reads were filtered via Bedtools. Peaks were called using MACS2 with flags “--broad --broad-cut-off 0.05 --keep-dup all”. We generated a consensus peak set per treatment with the overlapping peaks from the two biological replicates of each treatment. Differentially accessible chromatin analysis was made using the getDifferentialPeaks tool from HOMER, where a peak region with a fold change >1.5 and a Poisson p-value <0.0001 was considered to be differentially accessible. Signal plots were generated using the computeMatrix and plotProfile tools from Deeptools, peak annotation was made using the annotatePeaks tool from HOMER, plots were generated using the log CPM normalized counts calculated with the EdgeR package. Bigwig files for track visualization were generated using the bamCoverage tool from Deeptools. Public HUVEC H3K27me1 (ENCSR000AKL) and H3K27ac (ENCSR000ALB) bigwig and narrowPeak files were downloaded from ENCODE portal. The set of HUVEC enhancer regions was generated by filtering regions with overlapping H3K4me1, H3K27ac and ATAC-seq peaks with the intersect tool from Bedtools; track images were created using pyGenomeTracks. Transcription factor motif enrichment analysis in differentially accessible enhancers was made using the findMotifsGenome tool from the HOMER with “-size 50” flag to limit to ±50pb from the peak. Putative target genes for differential accessible enhancers were found using the Genomic Regions Enrichment of Annotations Tool (GREAT) with default parameters; these were an association rule of basal plus extension. This association rule assigns a ‘basal regulatory region’ 5 kb up- and 1 kb down-stream of the TSS, and each gene’s regulatory domain is then extended up to the basal regulatory region of the nearest upstream and downstream genes, but with a maximum extension of 1 Mb on each direction. Pathway enrichment analysis of putative target genes of DAEs was done using KEGG database within the EnrichR web tool with a p-value cut-off of 0.05.

### 4.10 NRF2 gene overexpression

The pHAGE-NFE2L2-IRES-EGFP plasmid (Addgene 116765) was used to produce lentiviral particles. Vector was also modified to have EGFP-only as negative control. Briefly, 8×10^6^ HEK293FT cells were seeded in a 15 cm dish with DMEM containing 10% FBS. The following day, cells were transfected using lipofectamine 3000 (Thermo Fisher Scientific) following manufacturer’s instructions: 20 μg of the transfer vector (with NFE2L2 or EGFP-only) and 10 μg of each packaging and envelope plasmids (psPAX2 and pMD2.G). Lentivirus-containing supernatant was collected at 48h and 72h post-transfection and then concentrated using Amicon Ultra 100 kDa filters (Merck, Rahway, NJ). Lentiviral titter was calculated by the % of GFP+ cells using flow cytometry on serial dilutions of concentrated lentivirus stock. Transduction of HUVEC was done by spinoculation were lentivirus supernatant was added to the medium (MOI=20) with 10 μg/mL polybrene and the plate with cells was centrifuged at 800 x g for 30 min at 32°C. Media was replaced after 24h. This approach yielded a 90% transduction efficiency (measured by % GFP+ via flow cytometry).

### 4.11 Real-time qPCR

HUVEC total RNA was isolated using QuickRNA Microprep kit (Zymo) following the manufacturer’s instructions. cDNA was synthesized from 500 ng of RNA with qScript cDNA SuperMix (Quantabio, Beverly, MA). RT-qPCRs were run on a CFX384 Real-Time Detection System (Bio-Rad, Hercules, CA). Reactions were prepared to a final 10 μL volume using 2X KAPA SYBR FAST qPCR Master Mix, primers (200 nM each), 25 ng of cDNA and nuclease-free water. Amplification program was: 95°C for 3 min, followed by 40 cycles of 95°C for 3 s and 60°C for 30 s, followed by a melt curve. Expression levels were calculated as relative to the housekeeping gene beta-actin. Primer sequences are shown in Table 3.

### 4.12 Statistical analysis

Data are presented as mean ± standard deviation; statistical tests used in each case are indicated in Figure legends, these tests were performed using either R or GraphPad Prism. Unpaired t-tests were used to compare the means of two different groups. The statistical significance cut-off used was a p-value <0.05 unless stated otherwise.

## Supplementary material

Summary of RNA-seq and ATAC-seq sequenced and aligned reads, genomic coordinates of differentially accessible enhancers and gene symbols of putative target genes and, tables of results of differential gene expression analyses and primers used.

## Data availability

Raw and processed sequencing data are available in the Gene Expression Omnibus repository under the accession number GSE241566

## Author contributions

MWV and VJV conceived and designed the study. MWV, BBG, OAG, JHG, NCD, MSN and CAA conducted the experiments. MWV, BBG, LGN and VJV analysed data; MWV performed the figures; TTL, SAR, LGN an VJV oversaw the study and funding. MWV and VJV drafted the manuscript; all authors critically revised the manuscript; all authors reviewed the manuscript and approved the submission.

## Acknowledgments

Marti Wilson-Verdugo conducted this study to fulfill the requirements of Programa de Doctorado en Ciencias Bioquímicas of Universidad Nacional Autónoma de México (UNAM), and received a doctoral scholarship from Consejo Nacional de Humanidades, Ciencias y Tecnologías (#CVU 853375). We thank to the UBM, UBMI, U de Computo and Bioterio from the IFC-UNAM. We thank Mayra Furlan-Magaril for critical reading of the manuscript.

---

**Supplemental figure S1.**
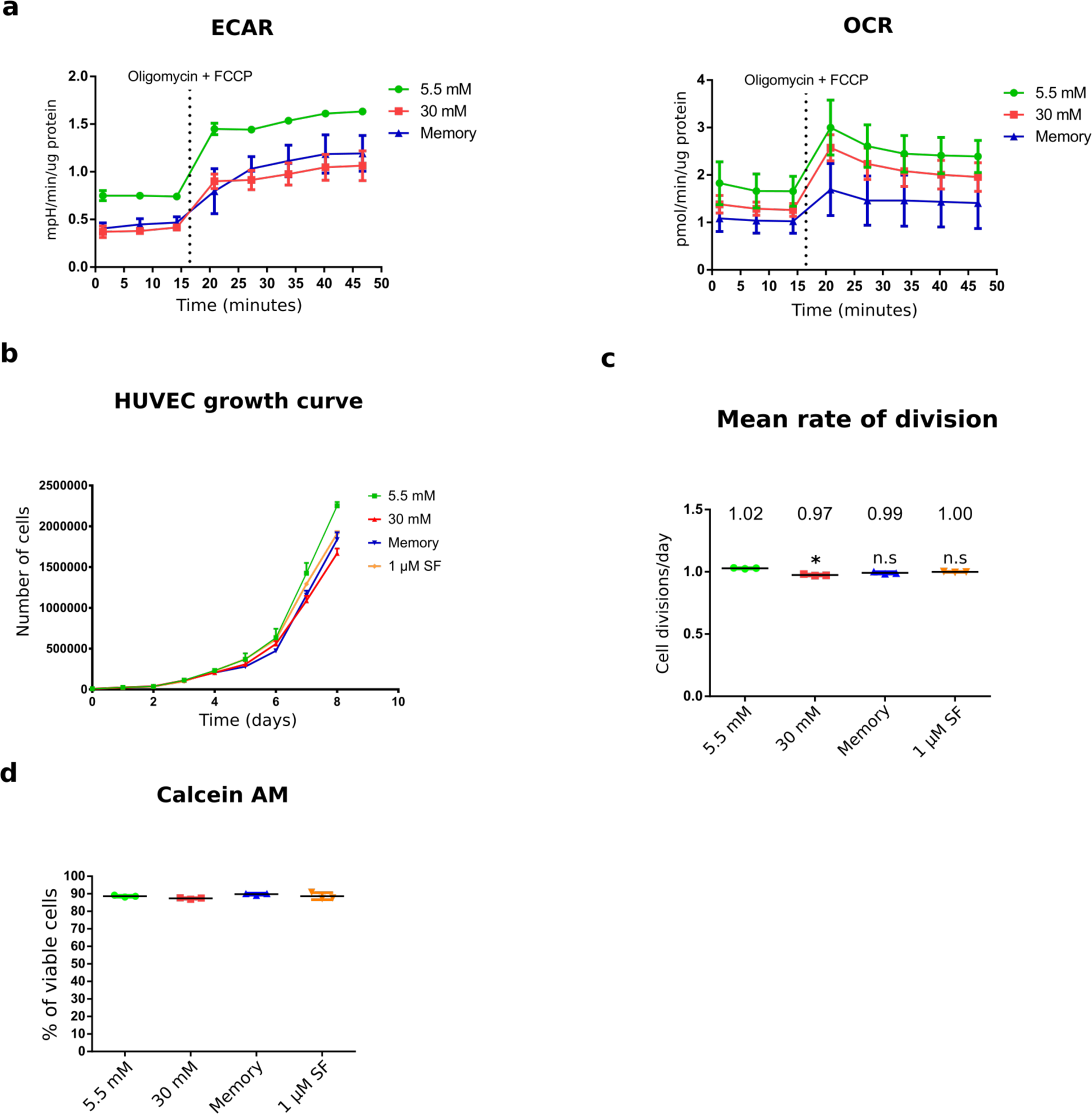
Metabolic phenotypes. **(a)** Time lapse of measurements of extracellular acidification rate (ECAR) and oxygen consumption rate (OCR) taken during the Seahorse XFp cell energy phenotype assay. Three baseline measurements were taken, followed by addition of oligomycin and FCCP to assess “stressed” ECAR and OCR measurements. Three biological replicates with three technical replicates each were used per condition. **(b)** 8 days growth curve of HUVEC in our control, high glucose, memory and 1 µM sulforaphane treatments. Cells were counted every day, biological triplicates with 2 technical replicates each were used. **(c)** Mean rate of cell divisions per day of HUVEC calculated at day 8 of culture. *P<0.05, n.s = not significant. **(d)** Calcein AM viability assay of HUVEC exposed to our treatments: control, high glucose, memory and 1 µM sulforaphane. This assay is based on conversion of non-fluorescent calcein AM to fluorescent calcein in viable cells. Differences against the control were examined by unpaired t-tests.

**Supplemental figure S2.**
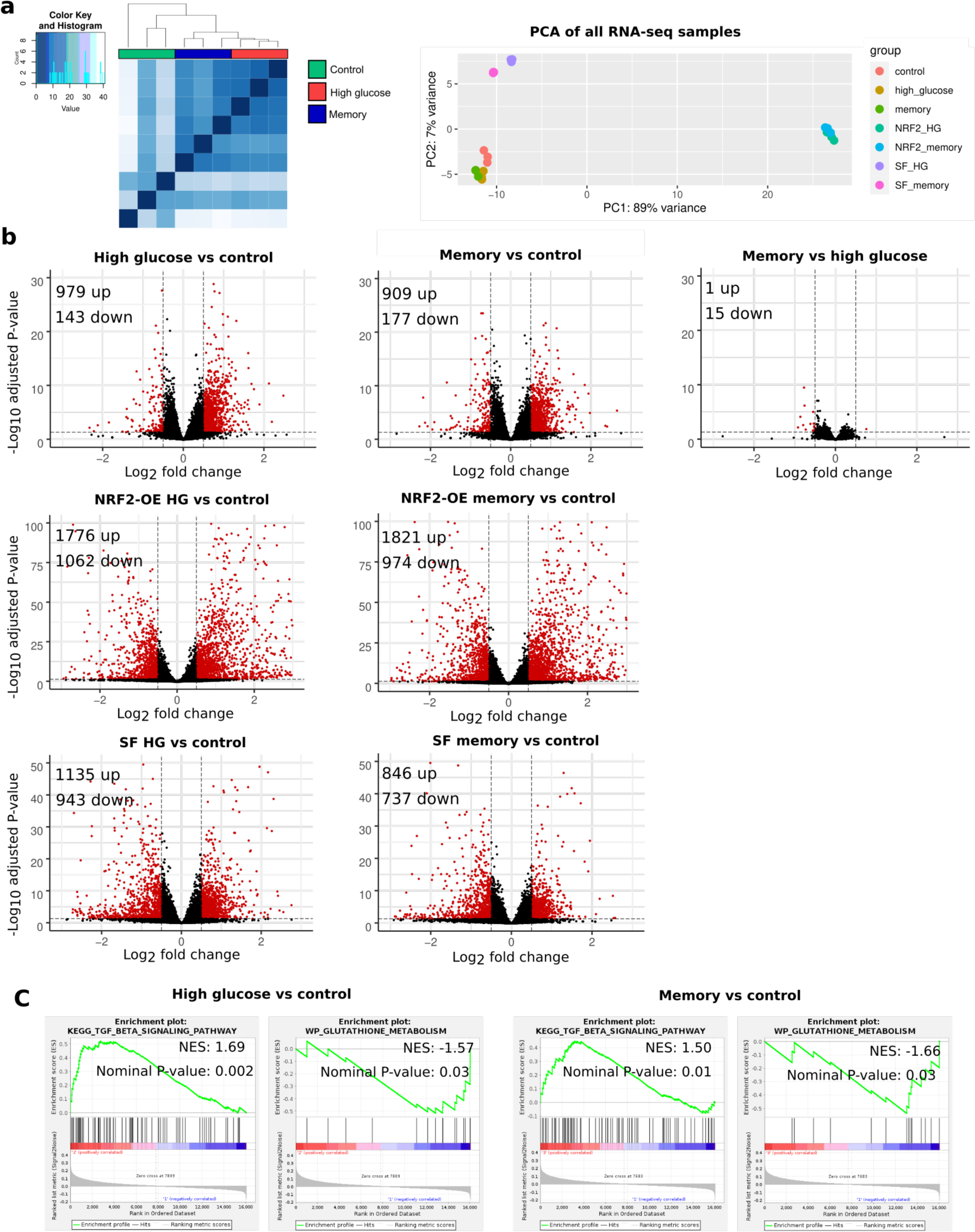
Transcriptional profiling. **(a)** Left: Clustering heatmap of control, high glucose (HG) and memory (M) RNA-seq samples. Euclidean distance was used as similarity measure. Right: Principal component analysis of all HUVEC RNA-seq samples used in this study. Three biological replicates were used per condition. **(b)** Volcano plots showing pairwise RNA-seq differential gene expression analyses between experimental conditions. Red dots are indicative of DEGs. The significance threshold used was an adjusted p-value < 0.05 and a log2 fold change >0.5 and <-0.5. **(c)** GSEA enrichment plots for the terms “TGF-beta signaling pathway (KEGG database)” and “Glutathione metabolism (Wikipathways database)” in two pairwise comparisons: high glucose versus control, and memory versus control. N.E.S; normalized enrichment score.

**Supplemental figure S3.**
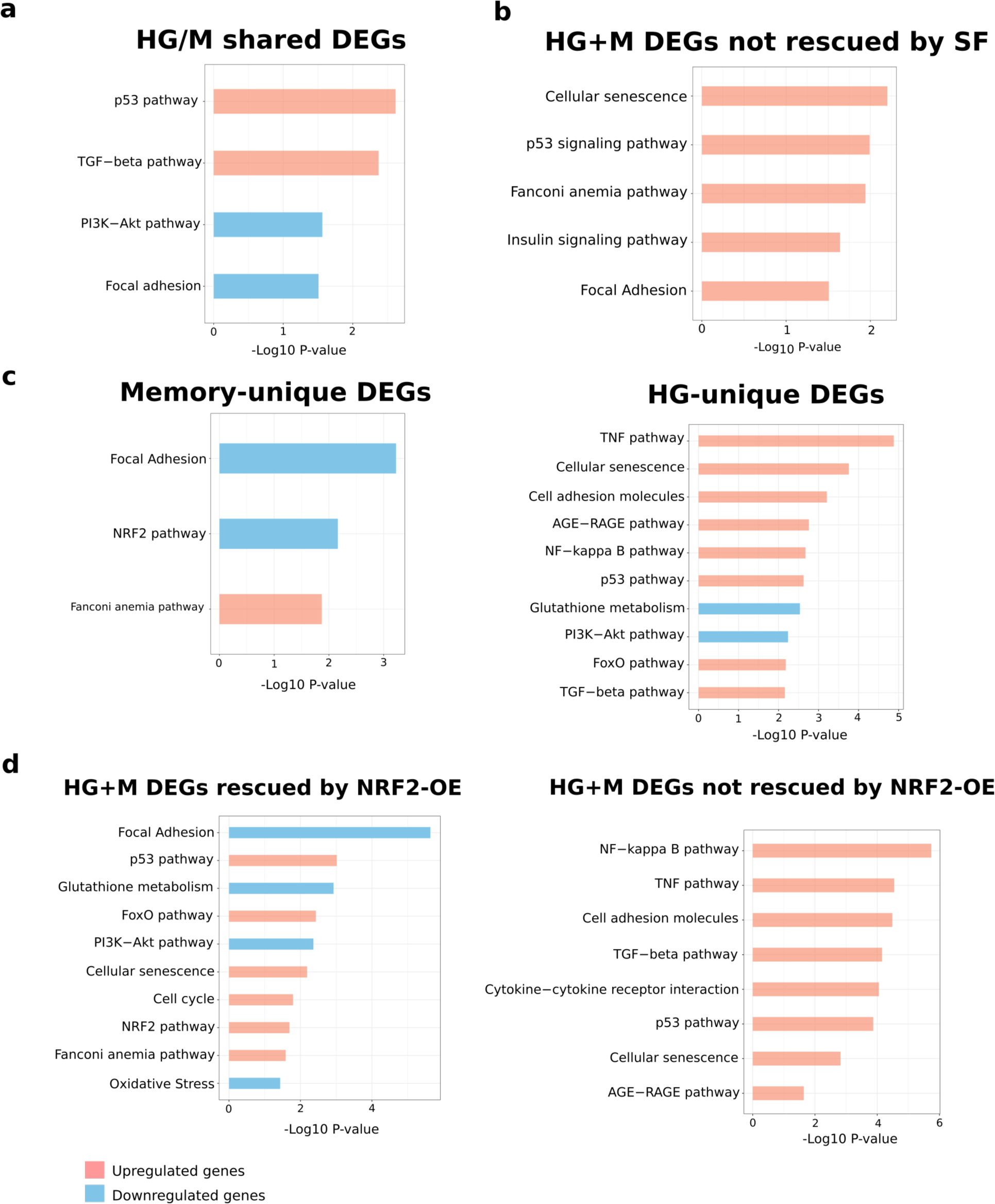
Pathway analysis of different subsets of high glucose and/or memory differentially expressed genes (DEGs). **(a)** Terms enriched in the subset of DEGs that are present in both HG vs. control and memory vs. control. (b) Terms enriched in HG+M DEGs not rescued by SF supplementation. (c) Terms enriched in DEGs unique to the memory vs. control subset (left) or the high glucose versus control subset (right). (d) Terms enriched in HG/M DEGs that were rescued (left) or not rescued (right) by NRF2 overexpression (OE).

**Supplemental figure S4.**
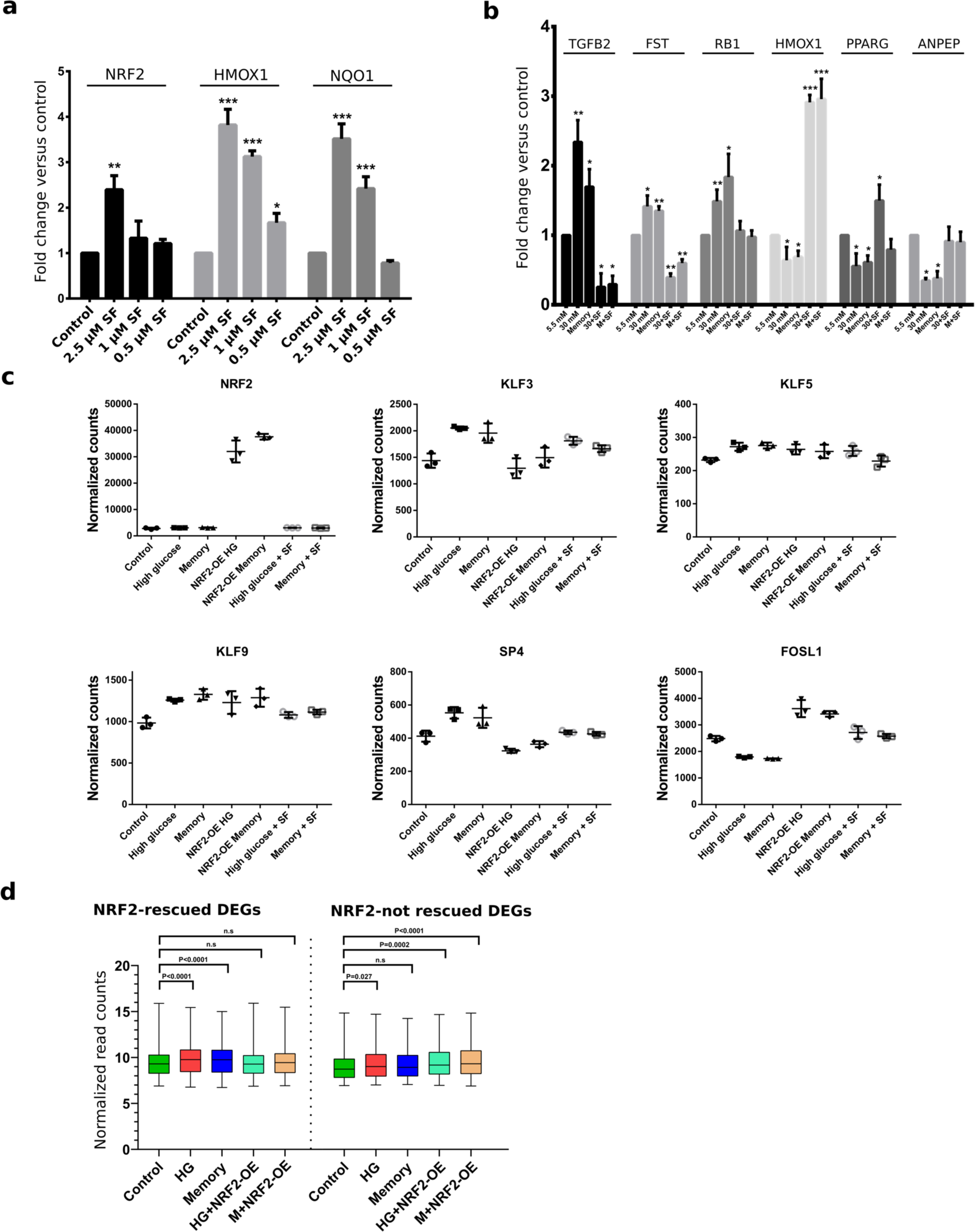
**(a)** RT-qPCR of NRF2 and its two target genes HMOX1 and NQO1 in HUVEC exposed to 0 (control), 2.5 μM, 1 μM and 0.5 μM sulforaphane at 5.5mM glucose. **(b)** RT-qPCR validation of six HG/M DEGs from our RNA-seq data. **(c)** DESeq2 normalized counts of NRF2 and transcription factors of the KLF/SP and bZIP families found to be differentially expressed in our high glucose and/or memory treatments. **(d)** Plot showing the changes in expression of HG/M DEGs after NRF2 OE, distinguishing between those rescued and not rescued by NRF2 OE. Differences against the control were examined by unpaired t-tests. *p<0.05, **p<0.01, ***p<0.001.

**Supplemental figure S5.**
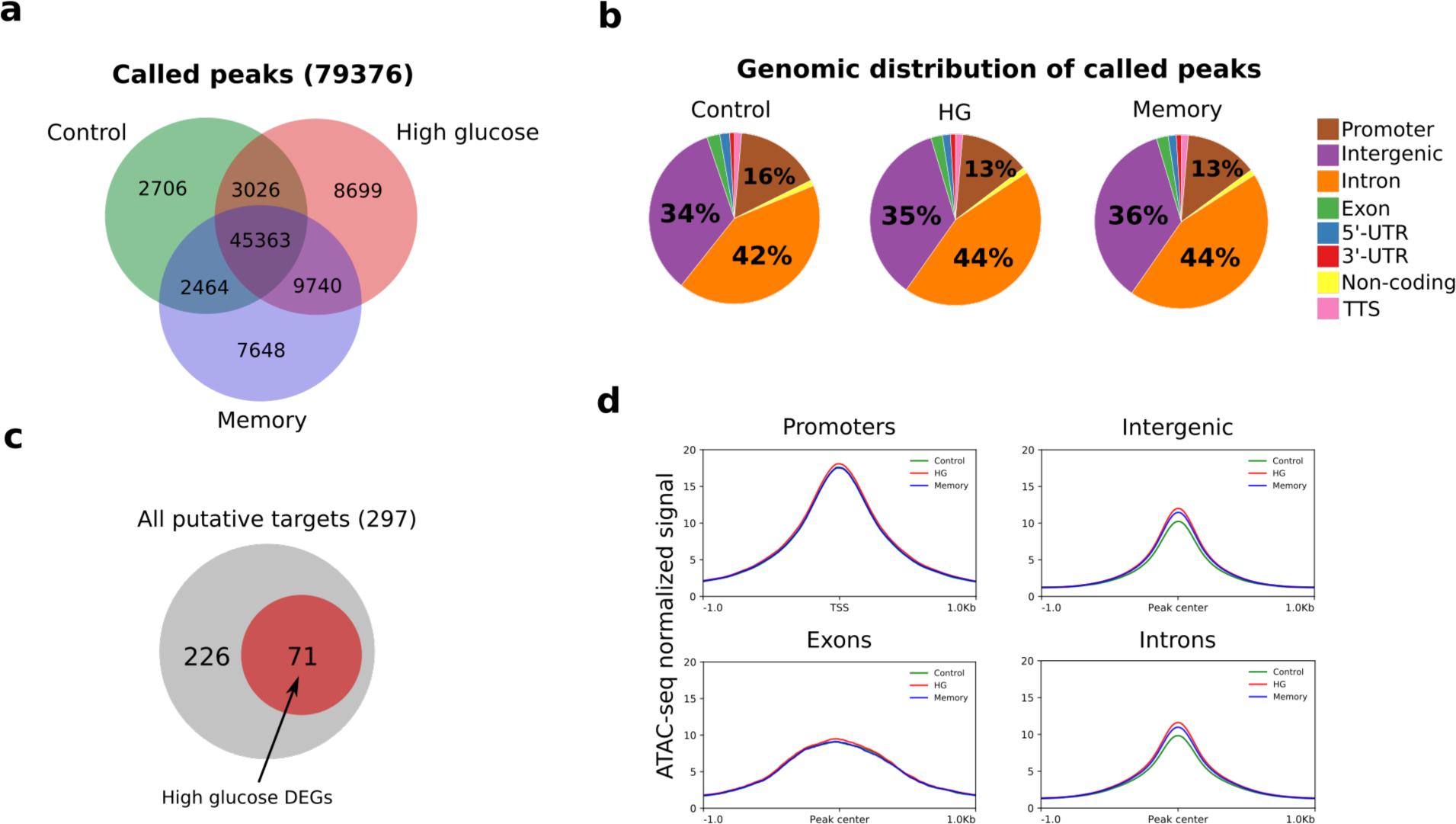
**(a)** Venn diagram showing the overlap of ATAC-seq called peaks between conditions. **(b)** Pie chart showing the percentage occupied by each genomic feature in the total number of ATAC-seq peaks called in each condition. **(c)** Diagram showing the proportion of HG/M DEGs that are putative targets of our HG/M DAEs according to the GREAT software. **(d)** ATAC-seq normalized signal in our control, HG and memory samples by genomic feature.

