## Supplemental tables for "NRF2 pathway activation reverts high-glucose-induced transcriptional memory in endothelial cells"

**Supplemental Table 1.** Summary of sequencing and mapping of RNA-seq samples.

| **Sample name** | **Sequenced pairs of reads (millions)** | **Mapped reads (millions)** |
| --- | --- | --- |
| Control A | 21.5 | 19 |
| Control B | 21.3 | 19.9 |
| Control C | 62.4 | 58.4 |
| High glucose A | 69.8 | 65.4 |
| High glucose B | 44.4 | 41.7 |
| High glucose C | 50.2 | 46.5 |
| Memory A | 57.4 | 52.9 |
| Memory B | 32.3 | 30 |
| Memory C | 48.5 | 44.9 |
| NRF2-OE HG_A | 26.3 | 24.6 |
| NRF2-OE HG_B | 16.7 | 15.6 |
| NRF2-OE HG_C | 14.3 | 13.3 |
| NRF2-OE Memory A | 39.2 | 36.8 |
| NRF2-OE Memory B | 20.4 | 18.6 |
| NRF2-OE Memory C | 27.9 | 25.6 |
| SF HG_A | 69 | 64.6 |
| SF HG_B | 20 | 18.4 |
| SF HG_C | 34.7 | 32.4 |
| SF memory A | 24.8 | 22.6 |
| SF memory B | 31.1 | 29 |
| SF memory C | 22.4 | 20.5 |

**Supplemental Table 2.** Summary of sequencing and mapping of ATAC-seq samples.

| **Sample name** | **Sequenced pairs of reads (millions)** | **Mapped reads (millions)** |
| --- | --- | --- |
| Control A | 39.5 | 37 |
| Control B | 51.5 | 49.3 |
| High glucose A | 40.1 | 38 |
| High glucose B | 38.4 | 36.6 |
| Memory A | 43.5 | 41.3 |
| Memory B | 37.2 | 35.6 |

**Supplemental Table 3.** List of primers used for RT-qPCR

| **Name (F is forward primer and R is reverse primer)** | **Sequence (5’-3’)** |
| --- | --- |
| TGFB2_F | CAGCACACTCGATATGGACCA |
| TGFB2_R | CCTCGGGCTCAGGATAGTCT |
| Actin_F | GCTATCCAGGCTGTGCTATC |
| Actin_R | TGAGGTAGTCAGTCAGGTCC |
| NQO1_F | GAAGAGCACTGATCGTACTGGC |
| NQO1_R | GGATACTGAAAGTTCGCAGGG |
| HMOX1_F | GACCCATGACACCAAGGACC |
| HMOX1_R | TCCACGGGGGCAGAATCTTG |
| PPARG_F | ACCAAAGTGCAATCAAAGTGGA |
| PPARG_R | ATGAGGGAGTTGGAAGGCTCT |
| RB1_F | TTGGATCACAGCGATACAAACTT |
| RB1_R | AGCGCACGCCAATAAAGACAT |
| ANPEP_F | TTCAACATCACGCTTATCCACC |
| ANPEP_R | AGTCGAACTCACTGACAATGAAG |
| FST_qF | TCTGCCAGTTCATGGAGGA |
| FST_qR | TCCTTGCTCAGTTCGGTCTT |
| NFE2L2_F | TCCAGTCAGAAACCAGTGGAT |
| NFE2L2_R | GAATGTCTGCGCCAAAAGCTG |
